# The potential of *Clonostachys rosea* as a biolarvicide against *Anopheles gambiae*

**DOI:** 10.64898/2025.12.29.695477

**Authors:** David T. Hayes, Patil Tawidian, Jennifer Phinney, Brandon L. Plattner, Kristin Michel

## Abstract

Mosquitoes vector multiple disease-causing pathogens detrimental to humans, imposing a high public health and economic burden. Chemical and biological larvicides are used in some settings to reduce mosquito populations as a supplement to adult population control. In this study, we assessed the larvicidal potential of a newly isolated strain of the hypocrealean fungus, *Clonostachys rosea*, by modifying a previously established laboratory-based experimental design. This design allowed us to distinguish between the larvicidal impact of the fungal culture supernatant and conidia. Our data show that *C. rosea* efficiently kills larvae of the African malaria mosquito, *Anopheles gambiae*, at an LC50 of 7.78 x 10^6^ for culture supernatant and 7.13 x 10^7^ conidia/mL for washed conidia. Fungal culture supernatant retained over 95% of its activity upon autoclaving, suggesting a temperature stable non-proteinaceous mycotoxin is responsible for larvicidal activity in the supernatant. Conidia were readily ingested, remained in the gut lumen, and did not germinate or translocate to the ectoperitrophic space nor the hemocoel. UV and heat-killed conidia retained larvicidal activity, demonstrating that no active secretion of toxins, fungal growth, or infection is required for killing mosquito larvae. However, autoclaving conidia abolished larvicidal activity, suggesting that larvae may not be killed via a mechanical mechanism, such as occlusion. Instead, our data demonstrate that *C. rosea* conidia kill *An. gambiae* larvae via the passive release of intracellular mycotoxins and have potential as a fungal biolarvicide to control *An. gambiae* populations.

**Highlights:**

- *Clonostachys rosea* is a potent larvicide of *Anopheles gambiae* mosquitoes.
- Larvicidal activity from *C. rosea* culture supernatant is heat stable.
- *C. rosea* conidia are ingested but remain in the gut lumen and do not germinate.
- *C. rosea* conidia passively release toxins ultimately leading to larval death.

## 1. Introduction

The African malaria mosquito, *Anopheles gambiae*, is a dominant vector of the malaria parasite (Sinka et al. 2012), causing significant morbidity, mortality, and economic burden to communities in Africa (Mezieobi et al. 2025). Mosquito control using indoor residual spraying (IRS) and long-lasting insecticidal nets (LLIN’s) is a core pillar of malaria transmission reduction, but its effectiveness is decreasing due to rapidly developing chemical insecticide resistance, contributing to the stalling progress in malaria transmission reduction (Ranson and Lissenden 2016, Killeen et al. 2017, WHO 2025).

Larval source management (LSM) is a promising supplemental intervention to adult control and entails removal of mosquito breeding sites and application of chemical or biological larvicides (Fillinger and Lindsay 2011, WHO 2013, 2024). LSM reduces adult and larval mosquito populations and malaria parasite incidence when applied effectively (Shililu et al. 2007, Fillinger and Lindsay 2011, Tusting et al. 2013, Choi et al. 2019). Therefore, LSM is becoming increasingly recognized as a crucial pillar to reduce malaria transmission, especially in urban and arid regions (WHO 2022, Newby et al. 2025, Okumu et al. 2025). However, like adult insecticides, resistance to chemical and bacterial larvicides, but notably not *Bacillus thuringiensis israelensis,* has been recorded (Su et al. 2018, 2019a, 2019b, Morales et al. 2019, Silva-Filha et al. 2021, Lopez et al. 2024, 2025, Clifton and Lopez 2025). Several entomopathogenic fungi (EPF) are larvicidal (Cafarchia et al. 2022) and could be used with traditional chemical and bacterial larvicides to reduce the impact of insecticide resistance (Scholte et al. 2004, Perumal et al. 2024).

Although interest using EPF as alternative mosquito larvicides is growing, their exact mode of killing remains highly understudied. Most often, to infect terrestrial insects, entomopathogenic fungal conidia first attach to the cuticle, then germinate and penetrate the integument (Hong et al. 2024). The fungus then enters the hemocoel, switching from a filamentous state to a yeast-type propagation strategy called blastospores (Butt et al. 2016, Hong et al. 2024). Blastospores kill by depriving the host of nutrients and releasing mycotoxins (Wang and Wang 2017, Ma et al. 2024). While it has been suggested that EPF follow this pattern of infection in larvae (Ghosh et al. 2021), most studies have demonstrated that conidia cannot attach to the cuticle and instead must be consumed to be larvicidal (Butt et al. 2013, Greenfield et al. 2014, Bawin et al. 2016). Once consumed, the fungi either penetrate the gut epithelium (Sweeney 1975, Noskov et al. 2019, 2021, Bitencourt et al. 2022, 2023) or release mycotoxins that damage the epithelial tissue leading to larval death (Lacey et al. 1988, Butt et al. 2013, Bawin et al. 2016, Bitencourt et al. 2024). These studies have focused mostly on a few fungal species of *Beauveria* and *Metarhizium*, necessitating research into other potential EPF (Cafarchia et al. 2022).

*Clonostachys rosea* is traditionally used as a biocontrol agent of fungal phytopathogens in agriculture but has recently been shown to be insecticidal. *Clonostachys rosea* has two EPA-approved strains for use as a fungicide in broad agricultural applications (Sun et al. 2020, Shafik 2025, U.S. EPA 2025). Toledo et al. 2006 first described *C. rosea* as insecticidal against two hemipteran species. Since then, additional studies have identified *C. rosea* as an insecticidal agent across a broad selection of insect species (Vega et al. 2008, Anwar et al. 2018, Mahmoudi et al. 2018, Kushiyev et al. 2022, Tamta et al. 2022), including two species of mosquito larvae, *Anopheles stephensi* and *Culex quinquefasciatus* (Prabhu and Kumar 2008). This demonstrates *C. rosea*’s potential as a mosquito larvicide. However, how *C. rosea* kills insects is currently unclear.

To address this knowledge gap, this study assessed the impact of the exposure of third instar *An. gambiae* larvae to culture supernatant and conidia of a newly isolated *C. rosea* strain. We quantified their potency by modifying a previously established laboratory-based experimental design (Tawidian et al. 2023, Hayes et al. 2025). We then determined whether ingestion of *C. rosea* conidia causes histologic changes to the larval gut and whether an active infection is required for larvicidal activity.

## 2. Methods

### 2.1. Isolation and molecular identification of *C. rosea* from mosquito larvae

*Clonostachys rosea* was isolated from field collected *Ae. albopictus* larvae as previously described (Tawidian et al. 2023, Hayes et al. 2025). In brief, larvae were collected from one location (N 39° 11ʹ35.5ʺ, W 96° 34ʹ15.7ʺ) in Manhattan, KS, USA using seed germination paper (Anchor Paper Co., St Paul, MN, USA) within oviposition cups. Larvae were washed with phosphate -buffered saline six times and homogenized in 50 µL of sterile water. The homogenate was plated onto potato dextrose agar (PDA) supplemented with antibiotics (25 mg/mL chloramphenicol and 100 mg/mL ampicillin [Sigma-Aldrich, St. Louis, MO, USA]) to inhibit bacterial growth. Plates were incubated in the dark at room temperature for 5 days. Unique fungal morphotypes were isolated for molecular identification and propagation.

To molecularly identify the isolate, total genomic DNA was extracted from mycelia and conidia using the DNeasy PowerSoil Kit (MoBio Laboratory, Carlsbad, CA, USA), following manufacturer’s protocol. The internal transcribed spacer (ITS) region was then amplified using the forward primer ITS1f (5’-CTTGGTCATTTAGAGGAAGTAA-3’) (Gardes and Bruns 1993) and reverse primer ITS4 (5’-TCCTCCGCTTATTGATATGC-3’) (White et al. 1990) following protocol described in Ihrmark et al. 2012. PCR products were purified using Qiaquick PCR purification kit (Qiagen, Germany). Purified ITS amplicons were sequenced by Sanger sequencing (GeneWiz, South Plainfield, NJ, USA) and blasted to the NCBI nucleotide database and compared to similar sequences. The ITS region sequence was submitted to GenBank and assigned accession number OP692517.1.

### 2.2. Mosquito maintenance

*Anopheles gambiae* larvae were reared at 27°C, 80% relative humidity (RH), and a 12L:12D photoperiod cycle as previously described (An et al. 2011), with freshly hatched larvae fed on 20 g/L baker’s yeast (Fleischman’s Active Dry Yeast, Associated British Foods, London, UK). Thereafter, larvae were fed daily using a slurry of 6.6 g/L baker’s yeast and 13.6 g/L ground fish food (Tetramin Tropical Flakes, Tetra, Blacksburg, VA, USA). Pupae were transferred to adult emergence cages, and the emerging adults were provided with 8% fructose (Emprove Chemicals, Sigma-Aldrich, MO, USA) supplemented with 2.5 mM PABA (Sigma-Aldrich, St. Louis, MO, USA) *ad libitum*. Female mosquitoes were blood fed with heparinized horse blood (PlasVacc, Templeton, CA, USA) via an artificial glass feeding system (Scientific Glass shop, Kansas State University, Manhattan, KS, USA) with expanded parafilm used as a membrane.

### 2.3. Exposure bioassay

*Anopheles gambiae* larvae were exposed to fungal and control treatments using a laboratory-based exposure assay previously established (Tawidian et al. 2023, Hayes et al. 2025). Mosquitoes were exposed at 27°C, 80% RH, and a 12L:12D photoperiod cycle. Individual L3 *An. gambiae* larvae (5 days old) were placed into individual wells of a 24-well plate and 1 mL of treatment or control solutions were pipetted into each well. Larvae were fed daily with 20 µL of fish food and yeast slurry and monitored daily for death, survival, and pupation.

### 2.4. *Clonostachys rosea* propagation and preparation of conidia and culture supernatant treatments

To produce large quantities of conidia, we first inoculated 150 mm diameter PDA plates with *C. rosea* hyphae and conidia. PDA plates were incubated at 25°C in the dark for 14-17 days. Conidia and extracellular metabolites were collected by flooding plates with approximately 20-30 mL sterile Milli-Q water. The solution was filtered through two layers of cheese cloth (Dritz, Spartanburg, SC, USA) to allow conidia and extracellular metabolites to filter through but retain mycelia and agar fragments. The conidia concentration of this initial suspension was estimated using a hemocytometer. Fig. S1 provides an overview of the four different fungal conidia and supernatant treatments used in the experiments. (1) The concentration of the initial suspension was diluted to 10^8^ conidia/mL and used as the 10^8^ treatment. (2) To prepare supernatant treatments, the 10^8^ treatment was centrifuged at 2,500 g for 20 minutes and the culture supernatant was collected, reducing its conidia concentration to approximately 4 x 10^3^ conidia/mL (SUP treatment). (3) Part of the culture supernatant was then autoclaved at 121°C and 15 psi for 40 minutes to inactivate any heat-labile metabolites and kill any remaining conidia (ACSUP treatment). (4) A washed conidia suspension was prepared by washing the remaining conidial pellet three times in 20-30 mL sterile Milli-Q water and adjusting its final concentration to 10^8^ conidia/mL (Washed conidia treatment).

### 2.5. Conidia viability

Conidia viability was assessed via germination assays by plating 50 µl of conidia-containing treatments onto PDA supplemented with antibiotics. Plates were incubated at room temperature in the dark for 18-24 hours post inoculation. Plates were stained with lactophenol blue and a total of 300 germinated or non-germinated conidia per treatment were counted. Conidia were considered viable if the length of its germ tube was at least twice the length of the diameter of the conidium (Francisco et al. 2006). Live conidia treatments were only used if they had a mean germination percentage above 90%.

### 2.6 Assessment of lethal concentration 50 (LC50) and LC90 values

Larvae were exposed to descending dilutions of *C. rosea* washed conidia and culture supernatant (Fig. S2) using the exposure bioassay described in 2.3. Culture supernatant was collected from 10^8^ conidia/mL *C. rosea* by centrifugation. Because we do not know the compounds present in the culture supernatant, we referred to this supernatant as the 10^8^ supernatant, corresponding to the number of conidia present in the solution prior to removal. The 10^8^ supernatant was further diluted with sterile Milli-Q water to 10^7^, 7.5 x 10^6^, 5 x 10^6^, 2.5 x 10^6^, 10^6^, and 10^5^ supernatant, respectively. Larvae were also exposed to a concentration gradient of washed conidia from 2.5 x 10^8^, 10^8^, 7.5 x 10^7^, 5 x 10^7^, 2.5 x 10^7^, 10^7^, to 7.5 x 10^6^ conidia/mL, respectively. LC50 and LC90 values of larval survival to pupation were calculated independently for both washed conidia and culture supernatant using a logit model in R (v4.4.3) (R Core Team 2021), with concentration as a fixed effect. LC50 and LC90 values were derived from the fitted model using the MASS package in R (Venables and Ripley 2002). All R code has been deposited in GitHub (https://github.com/Davidhayes2205/DavidHayes.Clonstachysrosea.Anophelesgambiae.larvalexposures.git).

### 2.7 Impact of culture supernatant and washed conidia on larval development

To assess sublethal effects, we recorded daily larval development to pupae during the LC50 and LC90 experiments. We separately compared larval development time across the concentration gradients of culture supernatant and washed conidia treatments using a multiple linear model with concentration and trial as fixed effects in R (v4.4.3) (R Core Team 2021). Least squares means (LSMs) and standard errors (SEs) were calculated for each concentration using the emmeans package in R (Lenth 2025).

### 2.8. Preparation of killed conidia treatments

To evaluate whether it is necessary for conidia to be viable in order to kill mosquito larvae, *C. rosea* conidia were killed using three methods. (1) autoclaving at 121°C and 15 PSI for 40 minutes (AC treatment), (2) heating in a 55°C water bath for 60 minutes (Heat treatment), and exposure to 253.7 nm ultraviolet light for 60 minutes (UV treatment). Conidia in all groups (autoclave, heat, or UV treated) exhibited normal conidia morphology but no germination after 24h, assessed as described in 2.5. Differences in survival to pupation between the killed-conidia and control treatments were assessed using a conditional logistic regression model with fungal treatment as fixed effect and stratified by trial. Multiple comparisons were performed and p-values adjusted using Bonferroni’s correction.

### 2.9. Quantification of ingested conidia

To assess ingestion of conidia, 3^rd^ instar *An. gambiae* larvae were exposed to 10^8^ conidia/mL washed conidia and fed only fish food (Tetramin Tropical Flakes, 20 μL per day, 13.6 g/L). After 3 days, living larvae were collected and washed twice in sterile milli-Q water. The guts of larvae were dissected and pooled into groups of 5 per 100 µL 0.01% Triton X-100 (Fisher Scientific, Hampton, NH, USA). Pooled guts were macerated using a plastic pestle and the number of conidia was quantified using a hemocytometer. The homogenized gut pools were also assessed by light microcopy for the presence or absence of germinating conidia and imaged using a Zeiss Axio Imager.A1 (Carl Zeiss AG, Germany).

### 2.10. Histopathological analyses

Third instar larvae were exposed to 10^8^ conidia/mL of washed conidia or water control and fed 20 µL fish food daily for 3 days. Control larvae were kept in water for the same period. Live larvae were directly placed in 10% neutral buffered formalin fixative (Fisher Scientific, Hampton, NH, USA) for 24 hours and dehydrated following an ascending ethanol series: 70%, 80%, 95%, 95%, 100%, 100% with 15 minutes per step. Samples were washed in xylene for clearing. Dehydration and clearing were performed using a Tissue-Tek VIP**^®^** (Sakura Finetek USA, Inc., Torrance, CA, USA). Larvae were then embedded in paraffin wax. Longitudinal sections were cut to 4 µm thickness using a Leica RM2135 microtome (Leica Biosystems, Nußloch, Germany). Sections were stained with hematoxylin and eosin (H&E) and Periodic acid–Schiff (PAS). Larvae were visualized using a standard upright Olympus BX53 light microscope with 10x ocular (22OD) and UPlanFL N objectives at additional magnification 4x, 20x, 40x, or 60x. High resolution images of representative larvae were collected using a DP27 Olympus microscope mounted camera, and data were collected and processed using the Olympus CellSens Standard (v4.4) software.

## 3. Results

### 3.1. Morphological and molecular identification of *Clonostachys rosea*

A fungal morphotype was isolated from *Ae. albopictus* mosquito larvae collected from artificial containers in Manhattan, KS, USA. Filamentous growth of the colonies initially colored white near the center and off-white towards the edges three days post inoculation (Fig. 1A and B). Seven days post inoculation, colonies had white-yellow colored filamentous growth at the edge, and the centers of the colonies were a bright yellow on the surface of the agar and the reverse side. The agar color itself changed to a bright yellow (Fig. 1C and D) around seven days post inoculation. Verticillium-type (Fig. 1E) and penicillium-type (Fig. 1F) conidiophores were observed under optical microscopy, typical for *C. rosea* isolates (Schroers et al. 1999, Schroers 2001). Conidia were hyaline and curved ellipsoidal in shape, also typical for *C. rosea* (Schroers 2001).

**Figure 1.**
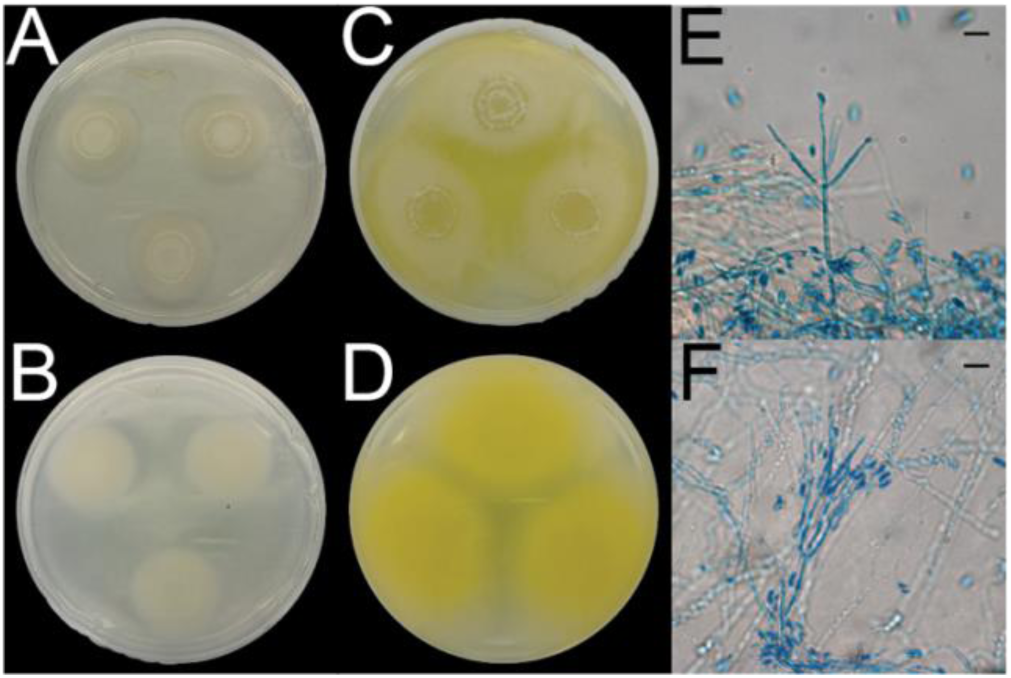
*Clonostachys rosea* colony and microscopic morphology. Left panel shows *C. rosea* growth three days post inoculation on the surface **(A)** and reverse side of PDA plates **(B).** Middle panel shows the top view **(C)** and bottom view **(D)** of the same colonies seven days post inoculation. Right panel shows the microscopic morphology of hyphae, conidia, and verticillium-type **(E)** and penicillium-type **(F)** conidiophores. Scale bar, 10 µm.

The ITS1, 5.8S, and ITS2 regions of the putative *C. rosea* isolate were sequenced to molecularly identify the morphotype. Comparison of the nucleotide sequence using the nonredundant GenBank database revealed >99.8% sequence identity to known *C. rosea* isolates, confirming the initial morphological species identification.

### 3.2. *Clonostachys rosea* conidia and culture supernatant are larvicidal

To determine whether *C. rosea* exhibited larvicidal activity against *An. gambiae*, we exposed 3^rd^ instar larvae to *C. rosea* conidia and culture supernatant and evaluated the percentage of larvae that pupated (Fig. 2A, Table S1). In the water-control treatment group, 89.8 % of larvae survived and pupated. When larvae were exposed to a combination of conidia and supernatant (10^8^ treatment), we observed 0% survival to the pupal stage, demonstrating a potent larvicidal activity of *C. rosea*. The larvicidal activity was fully retained in the culture supernatant (SUP), which by itself killed all exposed larvae, and 96.5% of its larvicidal activity remained after autoclaving (ACSUP). Furthermore, *C. rosea* conidia, washed to remove any culture supernatant also exhibited strong larvicidal activity, killing 91.7 % of all exposed larvae.

**Figure 2.**
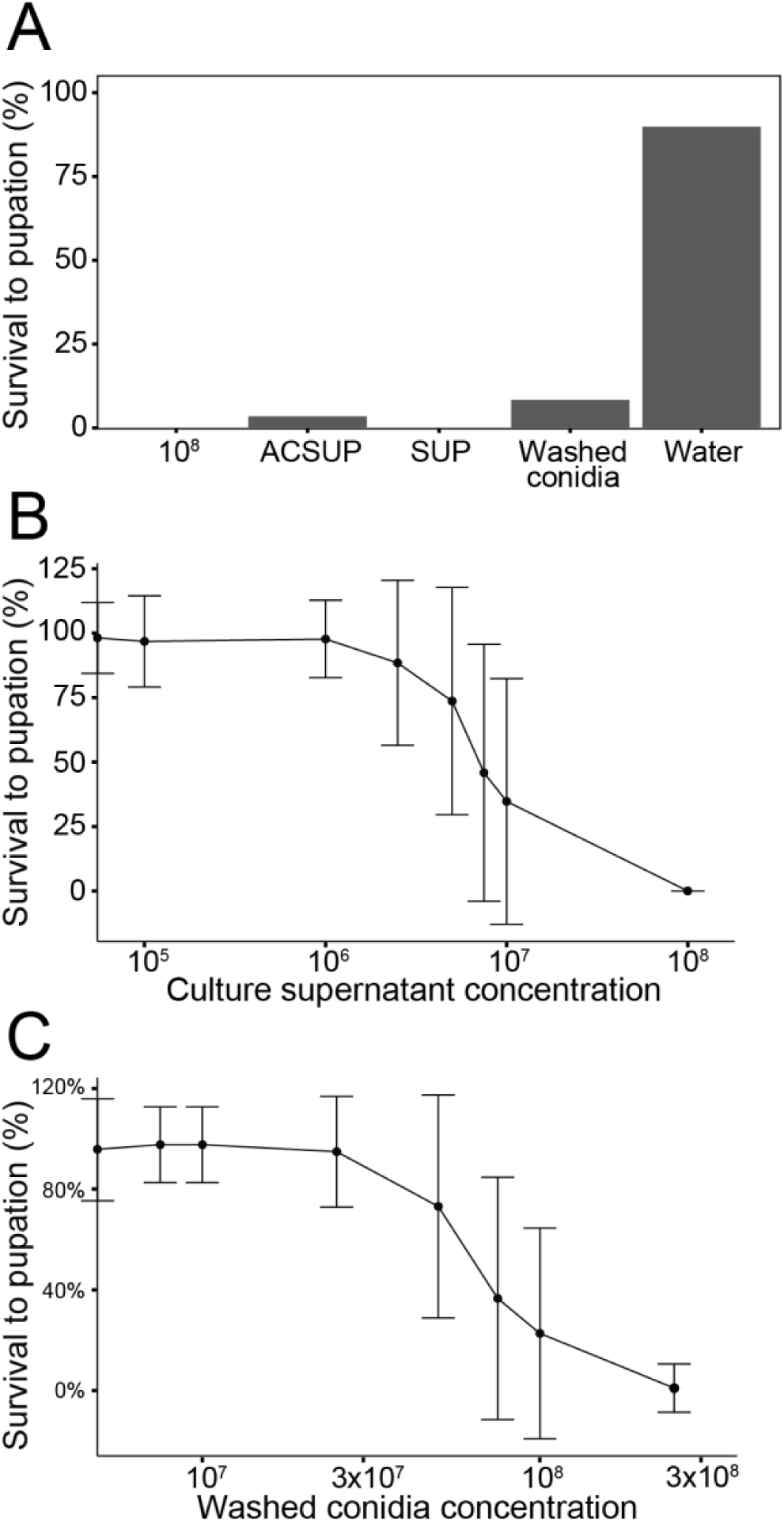
Survival to pupation of *An. gambiae* larvae decreased with increasing concentration of *C. rosea* culture supernatant treatments and washed conidia. **(A)** shows the percent survival to pupation of L3 *An. gambiae* larvae after exposure to washed conidia and culture supernatant treatments. The supernatant (SUP) treatment was prepared by centrifugating the 10^8^ treatment and was further autoclaved (ACSUP). The washed conidia treatment was washed 3x in water. The bottom two graphs show the dose-response curves of larvae exposed to descending concentrations of conidia-free culture supernatant **(B)** and washed conidia **(C)**. Culture supernatant concentrations refer to the number of conidia isolated from the same culture. Error bars depict Mean ± SD.

To quantify the potency of both culture supernatant and washed conidia we exposed larvae to descending concentrations and estimated LC50 and LC90 values. The LC50 and LC90 values for larvae exposed to culture supernatant were 7.78 x 10^6^, and 1.29 x 10^7^, respectively (Table 1, Table S1, Fig. 2B). The LC50 and LC90 values for larvae exposed to washed conidia were 7.13 x 10^7^ and 1.15 x 10^8^ conidia/mL (Table 1, Table S1, Fig. 2C).

**Table 1.**
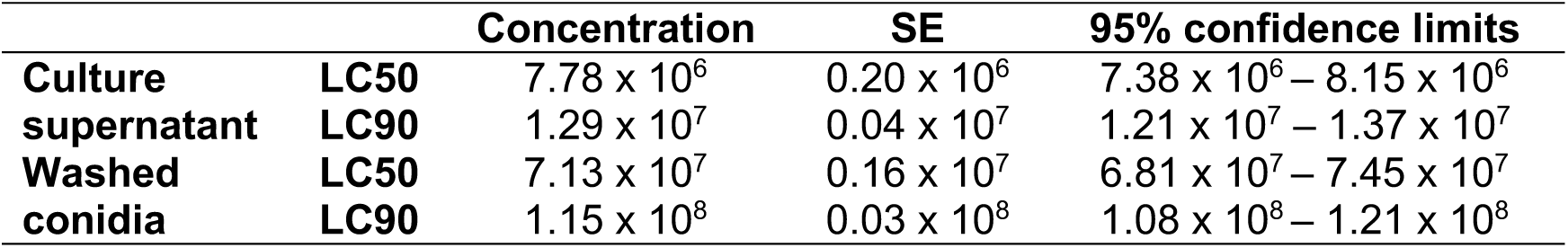
Lethal concentration 50 and LC90 estimations for *C. rosea* culture supernatant or washed conidia against *An. gambiae* larvae.

### 3.3. Sublethal doses of *C. rosea* conidia and culture supernatant increase development time to pupation

To evaluate whether *C. rosea* treatments had sublethal impacts to *An. gambiae* development, we quantified the time to pupation of larvae exposed to descending concentrations of culture supernatant and washed conidia (Table S1). As the concentration of culture supernatant and washed conidia increased, the time to pupation likewise increased (Table S2, Fig. 3). All culture supernatant concentrations (except for the 10^5^ supernatant dose) significantly increased time to pupation compared to the water control (ranging from 0.5 – 1.8 day delay, *P* < 0.01) (Figure 3A). Similarly, exposure to washed conidia increased time to pupation at sublethal doses but only when the concentration was 5 x 10^7^ conidia/mL or higher (0.6 – 1.8 day delay, *P* < 0.01) (Figure 3B).

**Figure 3.**
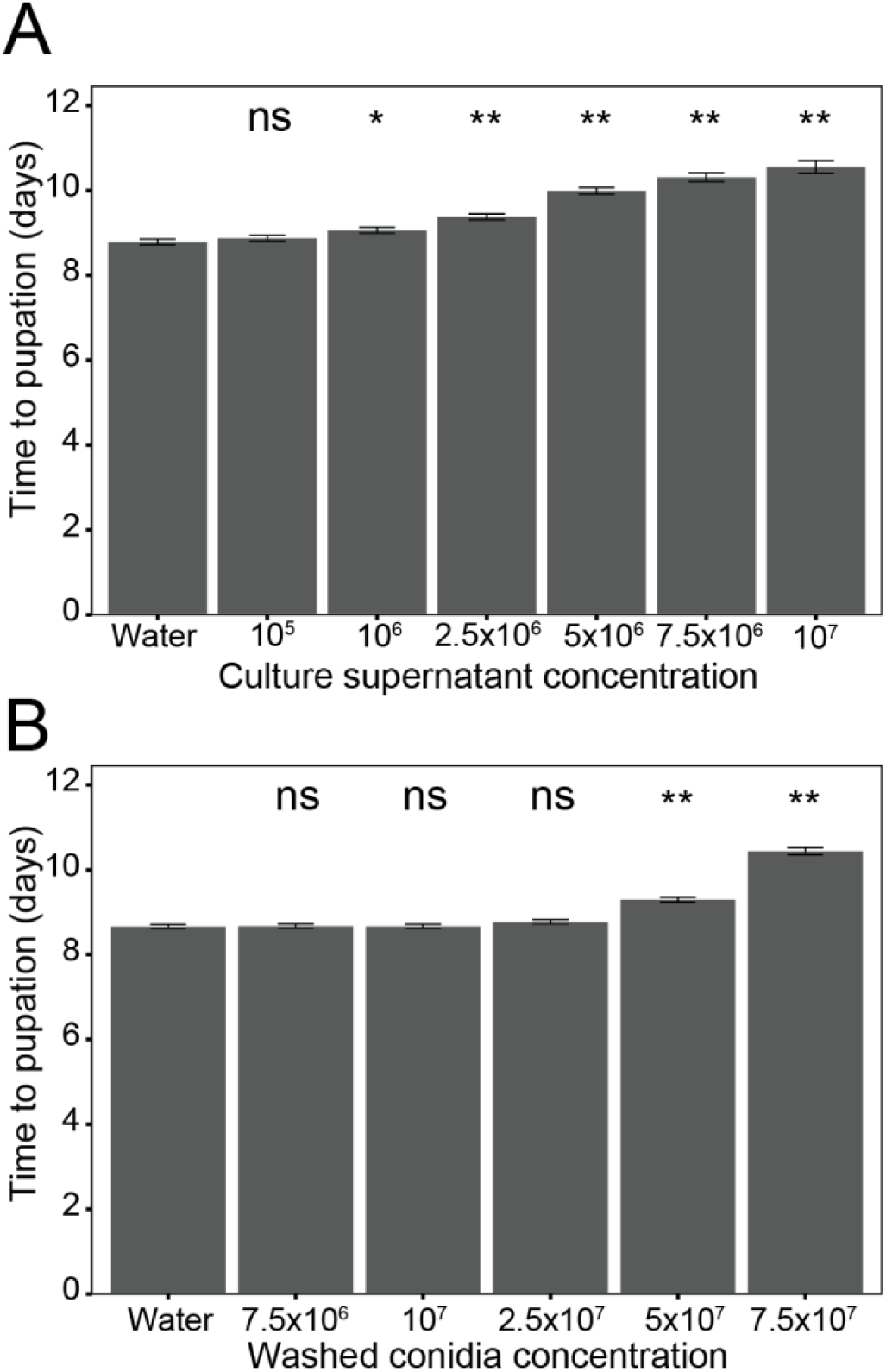
Time to pupation of *An. gambiae* larvae is extended after exposure to *C. rosea*. Increasing concentrations of *C. rosea* conidia-free culture supernatant **(A)** and washed conidia **(B)** extended significantly time to pupation of larvae exposed as L3 as compared to water-exposed controls. Results of the linear model are shown in Table S2. Conidia-free culture supernatant concentrations refer to the number of conidia isolated from the same culture. Data are shown as LSM ± SE. ns, not significant; *, *P*<0.01; **, *P*<0.0001.

### 3.4. Conidia are consumed into the gut but do not germinate or translocate into the hemocoel

Because *C. rosea* conidia exhibited potent larvicidal activity against *An. gambiae*, we further aimed to describe the general mode by which conidia killed larvae. Larvae were first exposed to 10^8^ conidia/mL washed conidia or water for 3 days. To formally demonstrate that conidia were ingested by larvae upon exposure, we dissected midguts to generate homogenates and prepared whole larvae for histologic analyses. Gut homogenates of *C. rosea* exposed larvae contained abundant conidia, typical in morphology to *C. rosea* (Fig. S3A), which were absent in the water-exposed control guts (Fig. S3B). We estimated a mean of 5.38 x 10^5^ conidia/gut (Fig. S4), demonstrating that larvae did consistently consume a high number of conidia in our experimental setup. Longitudinal sections of larvae exposed to *C. rosea* showed that the midgut lumen was filled with tightly packed conidia and few additional food particles (Fig. 4A-C). In contrast, the midgut lumen of water-exposed control larvae was filled with fish food particles of varied morphology (Fig. 4D-F). Although we observed high numbers of conidia within each gut, we did not observe germinating conidia or hyphae in either the midgut homogenates or longitudinal sections (Fig. S3A and Fig. 4A-C). In addition, fungal propagules were not observed in the ectoperitrophic space or in the hemocoel (Fig. 4A-C).

**Figure 4.**
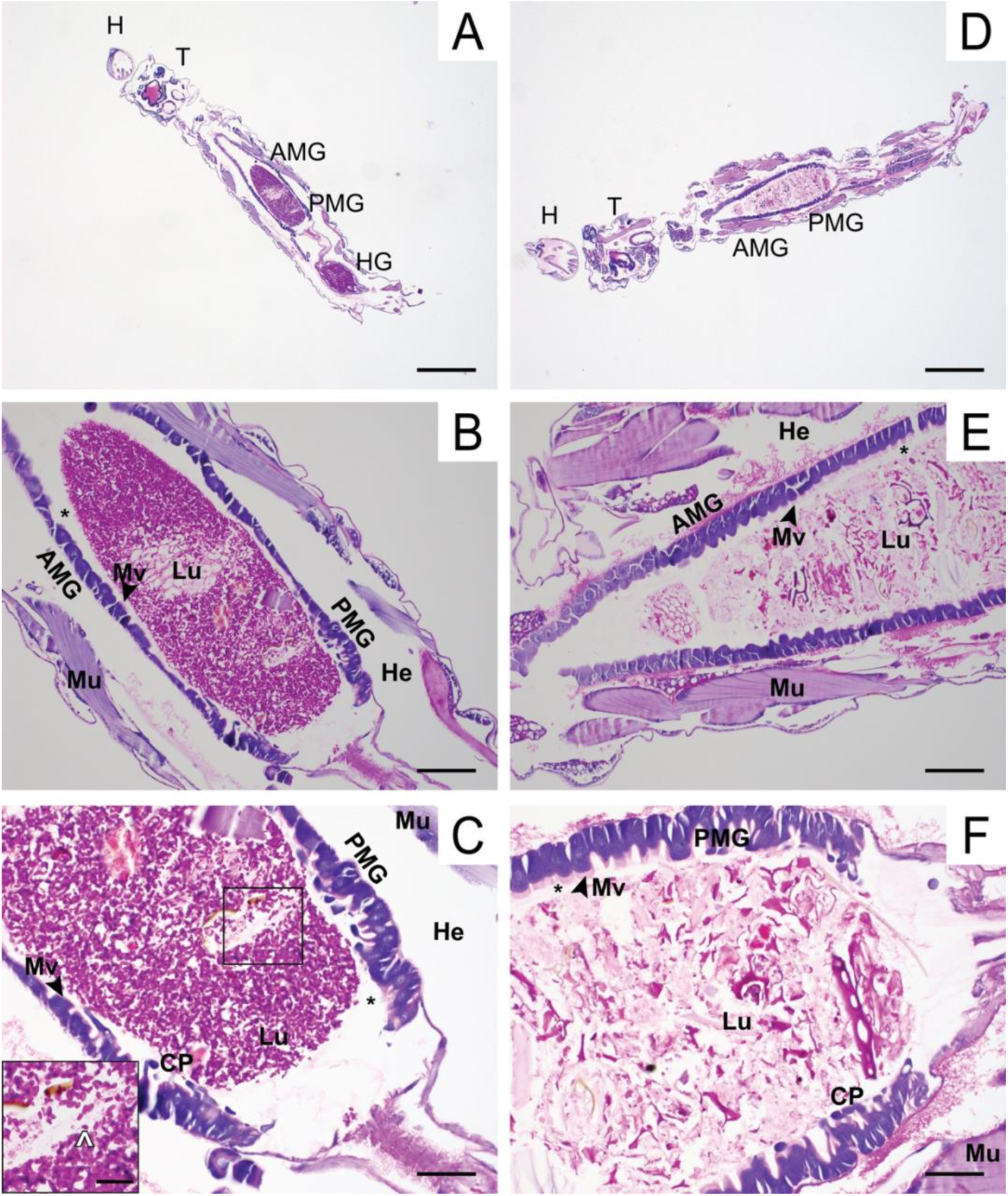
Conidia are contained within the midgut of *An. gambiae* larvae. Images show PAS-stained longitudinal sections of *Anopheles gambiae* larvae exposed for three days to *C. rosea* conidia **(A-C)** and water control **(D-F)**. Conidia are stained in dark magenta and show typical morphology for *C. rosea* **(inset, C)**. Epithelial cells are stained in dark purple. AMG, anterior midgut; CP, cytoplasmic projections; H, head; He, hemocoel; HG, hindgut; Lu, midgut lumen; Mu, muscle; Mv, microvilli; PMG, posterior midgut; T, thorax. Black asterisk, ectoperitrophic space; Black square, corresponding area of inset; White caret, an isolated conidium. Scale bars, 500 μm **(A, D)**, 100 μm **(B, E)**, 50 μm **(C, F),** 20 μm **(inset, C)**.

### 3.5. Histologic analysis of larval gut following conidia exposure

We further evaluated the midgut sections for histologic changes to the midgut epithelium and peritrophic matrix in the presence of conidia. In both control and treated larvae, the anterior midgut exhibited uniform and cuboidal shaped enterocytes with a thin surface layer of microvilli (Fig. 4B and E). The posterior midgut of larvae from both treatments were long and columnar shaped with cytoplasmic projections (also referred to as extracellular material) protruding from the enterocytes, and thick microvilli layers (Fig. 4C and F and Fig. S5). Lastly, the peritrophic matrix was observable and intact in both treatment groups (Fig. S5). Neither the midgut epithelium nor the peritrophic matrix exhibited obvious morphological differences in larvae exposed to *C. rosea* conidia compared to those exposed to water only.

In some larval sections we also observed either one or more gastric caeca or the hindgut in PAS-stained sections (Figs. S6 and S7). In one larva, the gastric caecum was largely devoid of purple staining or conidia-shaped propagules (Fig. S6A). In another larva, the lumen of the gastric caecum contained purple staining towards the periphery and contained a few conidia-shaped propagules (Fig. S6B). In addition, in one larva, the hindgut was distended and contained many conidia (Fig. S7A) and occasional fungal propagules were grouped in chains (Fig. S7B), potentially indicating growth within the hindgut lumen. In all histologic sections we examined, we did not find convincing evidence of penetration of epithelia nor translocation to or growth of *C. rosea* within the hemocoel.

### 3.6. Viable conidia are not necessary for larvicidal activity

Since neither gut homogenates nor histologic sections provided any evidence for an active infection, larval mortality could be caused by toxins that are either released passively (i.e., through digestion) or actively (i.e., secretion from conidia), or through a mechanical mechanism such as occlusion of the alimentary canal. To distinguish between these three processes, we exposed larvae to three killed-conidia treatments (autoclaved, heat, or UV-treated) using live washed conidia as positive and water as negative controls and assessed daily mortality (Table S1). 96.9% of larvae exposed to the AC treatment survived to pupa, similar to the 95.8% survival after water-control exposure (*P* > 0.999, Fig. 5), indicating that mosquito guts were not occluded by conidia. Larvae exposed to the heat-treated or UV-exposed conidia had significantly lower survival to pupation compared to the water control (heat: 24.4% survival, *P* < 0.0001; UV: 35.4% survival, *P* < 0.0001, compared to water-control, respectively). Heat-killed or UV-killed conidia were as effective in killing *An. gambiae* larvae as live washed conidia (*P* > 0.999, *P* = 0.776, Fig. 5), demonstrating that conidia viability was not required for larvicidal activity.

**Figure 5.**
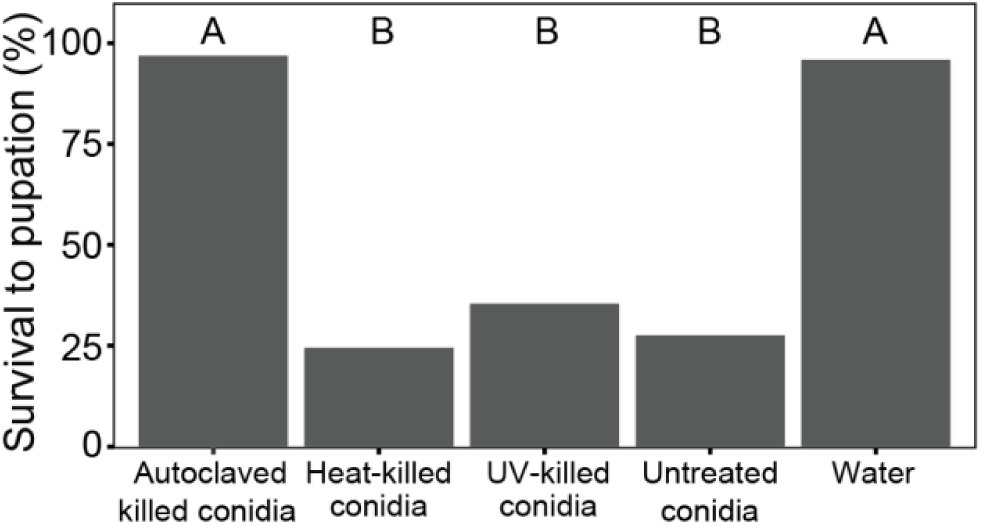
Conidia of *C. rosea* do not need to be viable to kill *An. gambiae* larvae. Graph shows percent survival to pupation of L3 larvae exposed to treated and untreated washed conidia. Treatment groups were statistically analyzed using conditional logistic regression model; different capital letters indicate statistically significant differences in larval survival to pupation.

## 4. Discussion

This study aimed to evaluate the larvicidal potential of a new isolate of *C. rosea* and describe its mode of killing of *An. gambiae* larvae. To do so, we modified a previously established laboratory based design (Tawidian et al. 2023, Hayes et al. 2025) to evaluate and describe the potency of *C. rosea* conidia and culture supernatant. Our data demonstrate that *C. rosea* conidia kill *An. gambiae* larvae following the passive release of toxins in the gut.

Both *C. rosea* conidia and culture supernatant were potent larvicides of *An. gambiae* larvae. Previous studies have shown *C. rosea,* and other *Clonostachys* species, with broad spectrum insecticidal activity (Sun et al. 2020, Reyes-Estebanez and Mendoza-de Gives 2025) including three species of mosquito larvae, *Aedes aegypti*, *An. stephensi*, and *Cx. quinquefasciatus* (Prabhu and Kumar 2008, Rodrigues et al. 2022). However, in these studies, either conidia and culture supernatant were not separated or the conidia collection method was not described, so it remained unclear what fungal culture components were the cause of the insecticidal activity. Our experimental design separated the culture supernatant from the conidia and demonstrated both to be larvicidal against *An. gambiae*.

In addition to killing, *C. rosea* conidia and culture supernatant delayed larval development to pupation when exposed to larvae at sublethal doses in a dose-dependent manner. Developmental delays are commonly observed for mosquito larvae exposed to sublethal doses of microbial- or chemical-based larvicides (Braga et al. 2004, Wang and Jaal 2005, Bara et al. 2014, Renuka et al. 2023, Tawidian et al. 2023, Hayes et al. 2025). To our knowledge, this is the first report describing the sublethal impact of *C. rosea* or other *Clonostachys* species towards insects. Developmental delays in insect larvae may be caused by damage to the intestinal epithelium and subsequently activated tissue repair, as shown in *Drosophila melanogaster* (Houtz et al. 2019, Nawrot-Esposito et al. 2020). Whether this mechanism contributes to the developmental delay caused by *C. rosea* remains unclear, as our histologic analyses so far did not find morphologic damage to midgut epithelia.

Our data strongly suggest that *C. rosea* conidia kill mosquito larvae via the passive release of mycotoxins following consumption, with several lines of supportive evidence. (i) In contrast to *Trichoderma asperellum,* which is suggested to penetrate the outer larval cuticle on *Anopheles* sp. (Ghosh et al. 2021), we did not find evidence of conidial attachment to the outer larval cuticle, nor did we find any fungal propagules in the hemolymph. Instead, our data are consistent with prior studies that show that fungal conidia in general are incapable of adhering to the larval cuticle, with the exception of the anal papillae or mouth parts, and instead must be consumed (Butt et al. 2013, Greenfield et al. 2014, Alkhaibari et al. 2016, Bawin et al. 2016). (ii) While *C. rosea* conidia were readily consumed, we also did not find any evidence of an active fungal infection via the gut after ingestion. *C. rosea* conidia filled the midgut lumen but did not germinate within the midgut or translocated from the gut lumen to the ectoperitrophic space or hemocoel. Indeed, few fungal species have been shown to cause active infection of mosquito larvae from the gut. *Culicinomyces sp.* and *Metarhizium spp.* conidia are able to form a penetration peg in the gut (Sweeney 1975, Bitencourt et al. 2022) and fungal propagules were found in the hemocoel after ingestion (Sweeney 1975, Noskov et al. 2019, 2021, Bitencourt et al. 2023). Similar to our observations of *C. rosea*, *Metarhizium anisopliae, Aspergillus clavatus, and B. bassiana* conidia do not germinate within the gut (Lacey et al. 1988, Butt et al. 2013, Bawin et al. 2016, Bitencourt et al. 2024). Instead, these fungal species are thought to kill mosquito larvae via the release of mycotoxins that damage midgut epithelial tissues. (iii) Heat-or UV-killed conidia retain larvicidal activity, demonstrating that viable conidia are not required to kill mosquito larvae, and instead strongly suggest the passive release of mycotoxins, likely due to conidia digestion.

The nature of these putative mycotoxins present in the culture supernatant and in the conidia are currently unknown. In this context it is interesting to note that the toxicity of the culture supernatant was largely insensitive to autoclaving, while the same treatment abolished larvicidal activity in conidia. This suggests that *C. rosea* produces several different molecules that are toxic to *An. gambiae* larvae. Many secondary metabolites of fungi, such as nonribosomal peptides and polyketides can induce apoptosis, which, if released in the gut, may lead to tissue damage (Trisciuoglio et al. 2008, Zhao et al. 2010, Ma et al. 2021, Toopaang et al. 2022). Interestingly, *C. rosea*’s genome putatively encodes for 32 polyketide synthase and 19 nonribosomal peptide synthetase genes (Karlsson et al. 2015), more than other larvicidal fungi such as *Metarhizium robertsii*, *M. anisopliae*, or *Beauveria bassiana* (Gao et al. 2011, Xiao et al. 2012). Therefore, it is possible that *C. rosea* synthesizes polyketides, nonribosomal peptides, or other autoclave-labile metabolites that are retained in conidia and released when ingested and digested by mosquito larvae.

Based on these data and prior studies (Butt et al. 2013, Bawin et al. 2016) we expected to find extensive tissue damage to the gut epithelium of *An. gambiae* larvae exposed to *C. rosea* conidia. To our surprise, we did not observe histologic evidence of damage to either the anterior or posterior midgut epithelium, nor the peritrophic matrix. One explanation may be the timing of our histologic analyses. Damage to the midgut epithelium may occur later than 3 days after exposure, as digestion of conidia may not have been sufficient at the time of collection. In addition or alternatively, damage leading to larval death may occur in tissues that we did not explicitly observe in our histological sections, such as the nervous system. Although, we did not observe damage to the midgut epithelium in our experimental design, other studies have shown that fungi are capable of killing by inducing caspase-mediated apoptosis in the midgut epithelium of mosquito larvae (Butt et al. 2013) and other insects (Kaczmarek et al. 2022, Kaczmarek and Boguś 2025). Future studies should focus on the identification of the mycotoxins responsible for larval mortality and their mode of action to elucidate how *C. rosea* kills mosquito larvae.

In conclusion, we report the identification of a mosquito-associated *C. rosea* isolate with larvicidal activity against *An. gambiae*. This isolate may be considered for future applications in mosquito larval control, because its activity is retained in killed conidia that cannot propagate upon release.

## Funding

KM is a member of the Vector-borne and Parasitic Disease team at Kansas State, which was supported by an Internal Gamechanging Research Initiation Program seed grant from the Office of the Vice President of Research. This study was supported by funding to KM from National Institutes of Health National Institute of Allergy and Infectious Diseases grant R01AI140760, and U.S. Department of Agriculture National Institute of Food and Agriculture Hatch projects 1021223 and 7007965. This is contribution 26-090-J from the Kansas Agricultural Experiment Station. The contents of this article are solely the responsibility of the authors and do not necessarily represent the official views of the funding agencies.

## Acknowledgements

We thank all members of the Michel laboratory for their help with mosquito rearing. The graphical abstract and Figures S1 & S2 were generated in BioRender.com.

## Author contributions (CRediT)

David Hayes: Conceptualization, Methodology, Validation, Formal analysis, Investigation, Data Curation, Writing – Original Draft, Writing – Review & Editing, Visualization.

Patil Tawidian: Conceptualization, Methodology.

Jennifer Phinney: Resources.

Brandon Plattner: Investigation, Resources, Data Curation, Writing – Review & Editing.

Kristin Michel: Conceptualization, Methodology, Validation, Resources, Data Curation, Writing – Original Draft, Writing – Review & Editing, Supervision, Project Administration, Funding acquisition.

## Notes

### Competing Interest Statement

The authors have declared no competing interest.

