## Supplementary information for "The potential of *Clonostachys rosea* as a biolarvicide against *Anopheles gambiae*"

**Supplementary Materials:**

**Supplementary Table S2** (Please note that Supplementary Table S1 is provided as a separate Excel file)

**Supplementary Figures S1-7**

**Table S2 – Time to pupation of *An. gambiae* larvae exposed to *C. rosea* culture supernatant and washed conidia**

| | Treatments | *N* | LSM[a] | SE[a] | Diff. to water control (*P*)[b] |
|---|---|---|---|---|---|
| Culture supernatant | Water-only | 211 | 6.78 | 0.068 | -- |
| | $10^5$ | 209 | 6.87 | 0.068 | 0.086 (0.370) |
| | $10^6$ | 199 | 7.06 | 0.070 | 0.279 (0.004) |
| | $2.5 \times 10^6$ | 184 | 7.38 | 0.073 | 0.593 (<0.0001) |
| | $5 \times 10^6$ | 158 | 7.99 | 0.079 | 1.204 (<0.0001) |
| | $7.5 \times 10^6$ | 101 | 8.31 | 0.100 | 1.528 (<0.0001) |
| | $10^7$ | 43 | 8.55 | 0.151 | 1.770 (<0.0001) |
| Washed conidia | Water-only | 206 | 6.66 | 0.052 | -- |
| | $7.5 \times 10^6$ | 210 | 6.67 | 0.051 | 0.010 (0.893) |
| | $10^7$ | 211 | 6.66 | 0.051 | 0.004 (0.950) |
| | $2.5 \times 10^7$ | 205 | 6.77 | 0.052 | 0.110 (0.131) |
| | $5 \times 10^7$ | 161 | 7.29 | 0.059 | 0.634 (<0.0001) |
| | $7.5 \times 10^7$ | 79 | 8.44 | 0.084 | 1.776 (<0.0001) |

[a]LSM: Least squares mean; SE: Standard error

[b]Compared to water-only control

**Supplementary figures with legends**

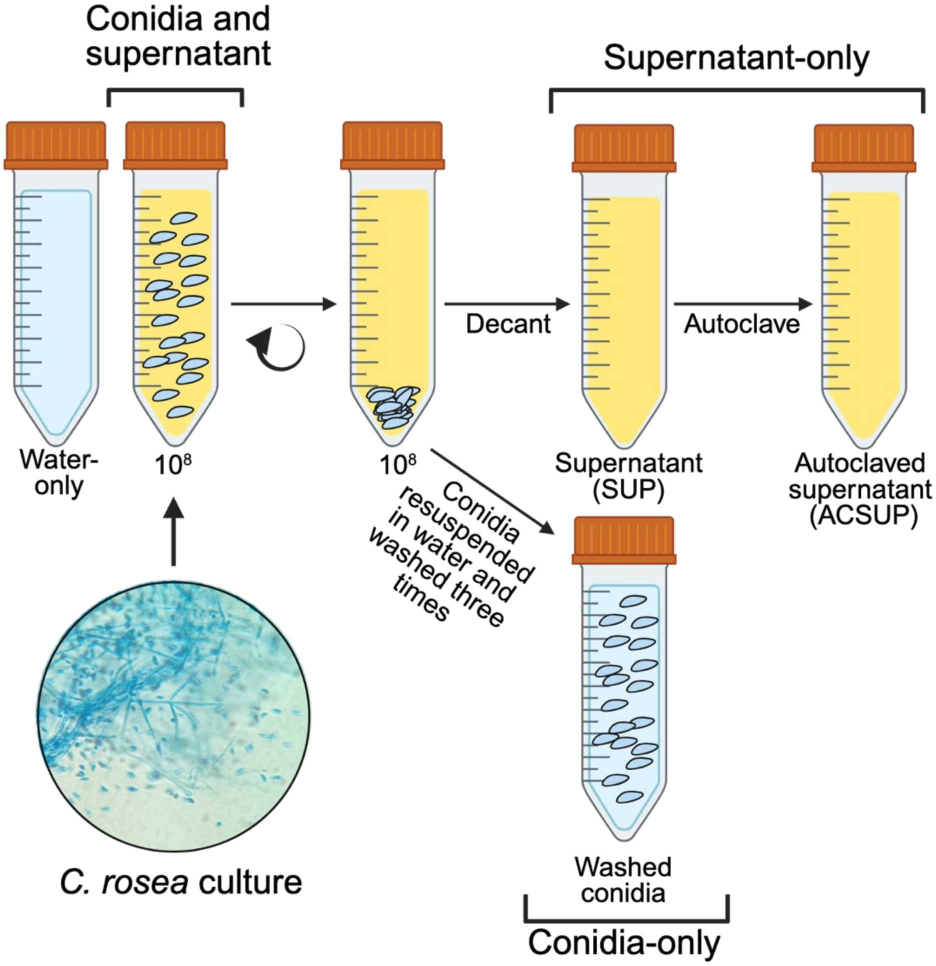


**Figure S1 – Graphical methods of *Clonostachys rosea* initial treatment preparations.**

Fungal conidia and culture supernatant were collected from potato dextrose agar to prepare treatments. Created in https://BioRender.com

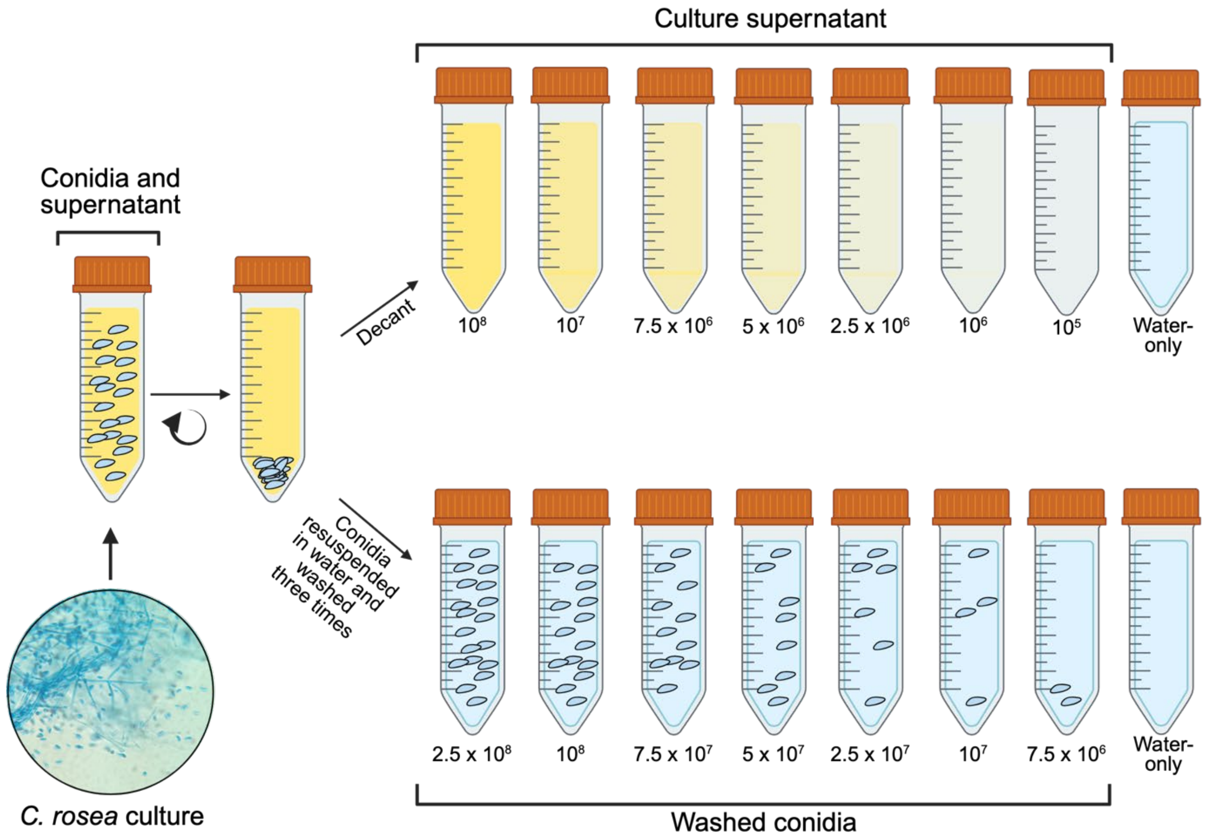


**Figure S2 – Graphical methods of *Clonostachys rosea* treatment preparations for LC50 and LC90 calculations.** Fungal conidia and culture supernatant were collected from potato dextrose agar to prepare treatments. Washed conidia and culture supernatant were diluted to prepare seven descending concentrations. Created in https://BioRender.com

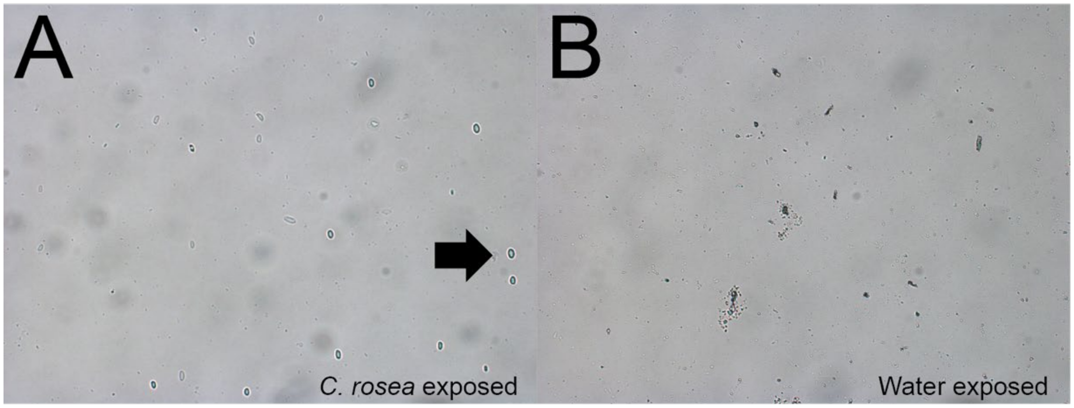


**Figure S3 – Representative images of *An. gambiae* larval gut homogenates.** Images of larval guts pooled after exposure to **(A)** *C. rosea* or **(B)** water for 3 days. Arrow points to putative *C. rosea* conidia based on morphology.

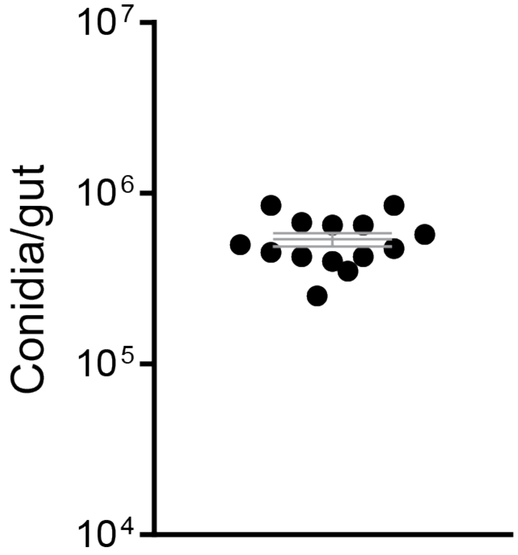


**Figure S4 – Quantity of *C. rosea* conidia consumed by *An. gambiae* larvae.** Number of conidia per gut of *An. gambiae* larvae after exposure to *C. rosea* for three days. Each dot represents the mean conidia per individual gut from a pool of 5 mosquito midguts and gastric caeca. Error bar depicts mean ± standard error

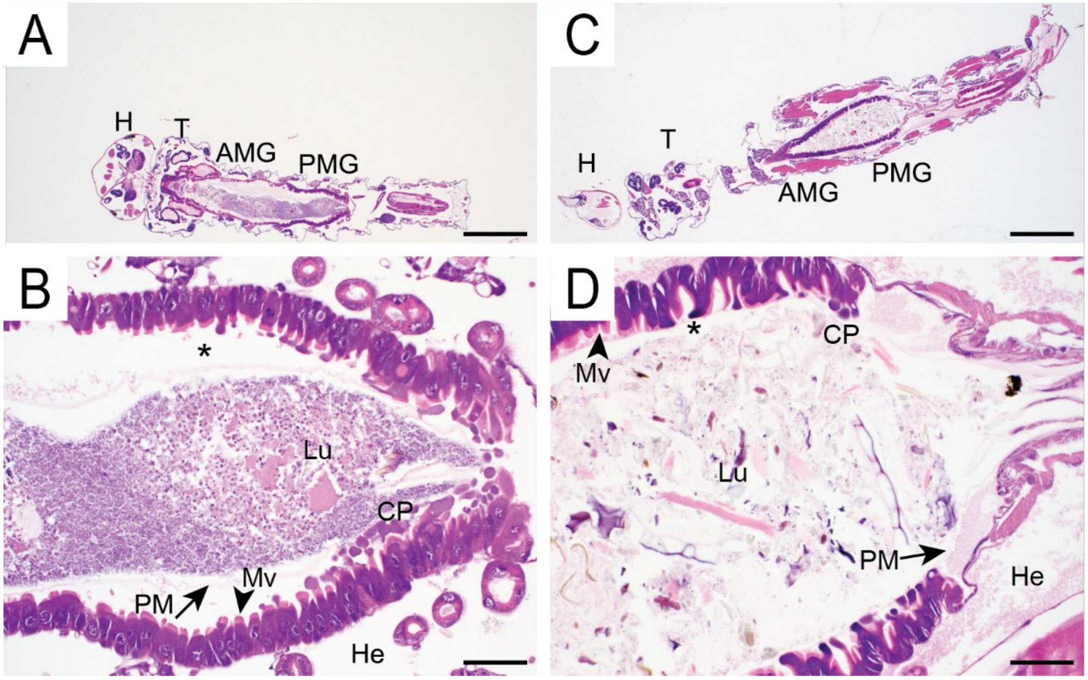


**Figure S5 – H&E-stained longitudinal sections of *An. gambiae* larvae.** *Anopheles gambiae* larvae exposed to *C. rosea* conidia **(A, B)** and *An. gambiae* larvae exposed to water control **(C, D)**. AMG, anterior midgut; CP, cytoplasmic projections; H, head; He, hemocoel; Lu, midgut lumen; Mv, microvilli; PM, peritrophic matrix; PMG, posterior midgut; T, thorax; asterisk, ectoperitrophic space. Scale bars, 500 μm **(A, C)** and 50 μm **(B, D).**

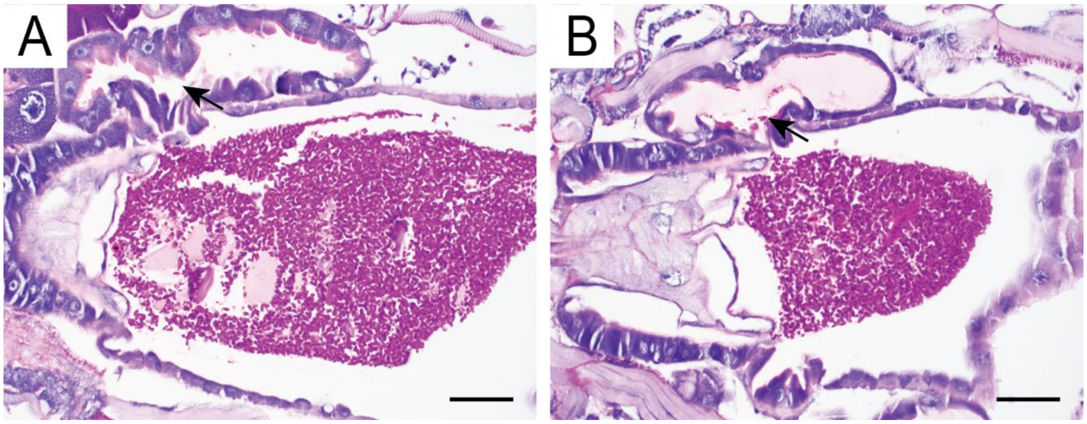


**Figure S6 – The presence and absence of conidia within gastric caeca of *An. gambiae* exposed to *C. rosea*.** Histologic sections of *An. gambiae* midguts, including a gastric caecum stained with PAS. Gastric caecum without intraluminal propagules **(A)** and with intraluminal propagules **(B)**. Arrows point to gastric caeca. Scale bar, 50 μm.

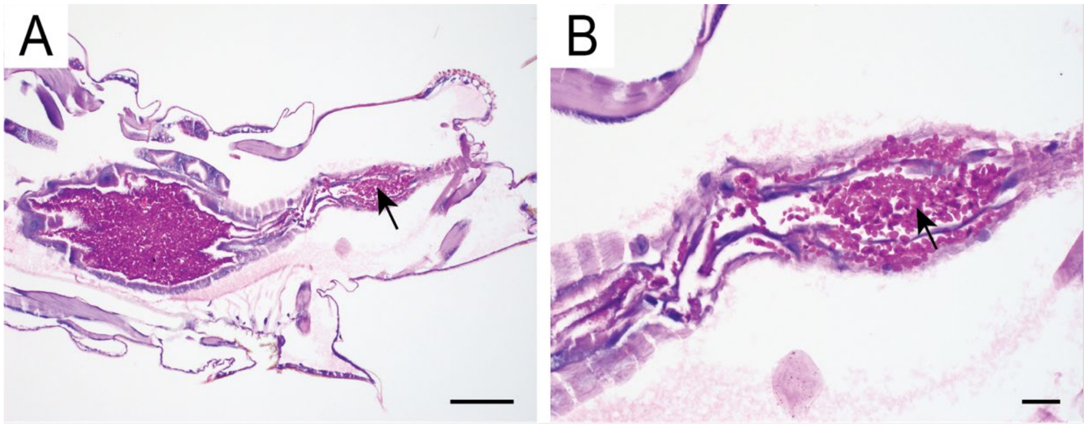


**Figure S7 – Histologic section of the hindgut of an *An. gambiae* larva exposed to *C. rosea* conidia.** Images of an *An. gambiae* hindgut filled with *C. rosea* conidia. Sections were stained with PAS. Scale bars, 100 μm **(A)** and 20 μm **(B).** Arrows point to *C. rosea* conidia.
